# Modeling the profibrotic microenvironment *in vitro*: model validation

**DOI:** 10.1101/2023.12.12.571237

**Authors:** Olga Grigorieva, Natalia Basalova, Uliana Dyachkova, Ekaterina Novoseletskaya, Maksim Vigovskii, Mikhail Arbatskiy, Maria Kulebyakina, Anastasia Efimenko

## Abstract

Establishing the molecular and cellular mechanisms of fibrosis requires the development of validated and reproducible models. The complexity of *in vivo* models challenges the monitoring of an individual cell fate, in some cases making it impossible. However, the set of factors affecting cells in *in vitro* culture systems differ significantly from *in vivo* conditions, insufficiently reproducing living systems. Thus, to model profibrotic conditions *in vitro*, usually the key profibrotic factor, transforming growth factor beta (TGFβ-1) is used as a single factor. TGFβ-1 stimulates the differentiation of fibroblasts into myofibroblasts, the main effector cells promoting the development and progression of fibrosis. However, except for soluble factors, the rigidity and composition of the extracellular matrix (ECM) play a critical role in the differentiation process. To develop the model of more complex profibrotic microenvironment *in vitro*, we used a combination of factors: decellularized ECM synthesized by human dermal fibroblasts in the presence of ascorbic acid if cultured as cell sheets and recombinant TGFβ-1 as a supplement. When culturing human mesenchymal stromal cells derived from adipose tissue (MSCs) under described conditions, we observed differentiation of MSCs into myofibroblasts due to increased number of cells with stress fibrils with alpha-smooth muscle actin (αSMA), and increased expression of myofibroblast marker genes such as collagen I, EDA-fibronectin and αSMA. Importantly, secretome of MSCs changed in these profibrotic microenvironment: the secretion of the profibrotic proteins SPARC and fibulin-2 increased, while the secretion of the antifibrotic hepatocyte growth factor (HGF) decreased. Analysis of transciptomic pattern of regulatory microRNAs in MSCs revealed 49 miRNAs with increased expression and 3 miRNAs with decreased expression under profibrotic stimuli. Bioinformatics analysis confirmed that at least 184 gene targets of the differently expressed miRNAs genes were associated with fibrosis. To further validate the developed model of profibrotic microenvironment, we cultured human dermal fibroblasts in these conditions and observed increased expression of fibroblast activation protein (FAPa) after 12 hours of cultivation as well as increased level of αSMA and higher number of αSMA+ stress fibrils after 72 hours. The data obtained allow us to conclude that the conditions formed by the combination of profibrotic ECM and TGFβ-1 provide a complex profibrotic microenvironment *in vitro*. Thus, this model can be applicable in studying the mechanism of fibrosis development, as well as for the development of antifibrotic therapy.

## Introduction

Fibrosis and fibrotic diseases are responsible for up to 50% of mortality in developed countries [1]. Approaches to the treatment of fibrosis are largely limited, despite the significant development of knowledge and mechanisms of this process that has accumulated recently. Thus, pirfenidone, one of the approved drugs for idiopathic pulmonary fibrosis (IPF) treatment, is considered to suppress the signaling pathway of the key profibrotic factor, transforming growth factor beta 1 (TGFβ-1). However, the use of pirfenidone does not stop the development of IPF, but only slows it down [2]. Importantly, the addition of TGFβ-1 to cultured cells is the most routine way to mimic a profibrotic environment and stimulate myofibroblast differentiation *in vitro* [3,4,5]. Considering the complexity of fibrosis and accepting that blocking of TGFβ-1 activity is not enough to suppress this process, we can conclude that the mentioned model based only on TGFβ-1 supplementation does not sufficiently reproduce the profibrotic conditions. The influence of microenvironment leads to differentiation of myofibroblasts which are the main effector cells promoting the development and progression of fibrosis. Myofibroblasts can differentiate from a wide range of stromal cells, including tissue fibroblasts, mesenchymal stromal cells, pericytes, circulating fibrocytes [6]. Activation of these cell types is characterized by the expression of the fibroblast activation protein (FAPa). Myofibroblasts are characterized by the secretion of large amounts of extracellular matrix (ECM) proteins and ability to contract ECM through the development of stress fibrils, including alpha-smooth muscle actin (αSMA), and mature focal junctions as well as fibronexuses [7].

ECM is considered as one of the crucial components that can independently initiate and maintain the development of fibrosis. Many studies demonstrate that the composition and stiffness of ECM determines the cell differentiation. Thus, an alternative splice form of fibronectin with an additional domain A (EDA-fibronectin) accumulates during the development of fibrosis and initiates the differentiation of new myofibroblasts [8]. Increased ECM stiffness leads to the enlargement of myofibroblast pool through the activation of Hippo signaling [9], release of ECM-anchored TGFβ-1 [10] as well as stimulation of the profibrotic regulatory microRNA expression [11]. These and other data confirm the exclusive role of ECM in stimulating the induction and progression of fibrosis. Regarding this case, the models used *in vitro* include either contact of fibroblasts with one of the components of ECM (collagen [12], fibronectin [13], hyaluronic acid [14], etc.), or imitate increased stiffness [15], thus not mimicking its multicomponent composition and architecture.

We have previously shown that decellularized ECM produced by human skin fibroblasts when grown in high density (so called “cell sheets”) with the addition of ascorbic acid, retains its native architecture and multicomponent composition, and has the ability to support the induced differentiation of committed progenitor cells [16]. It is known that TGFβ-1 induces fibroblast differentiation into myofibroblasts and stimulates the production of ECM proteins including type I and III collagen, EDA-fibronectin [4]. In this work, we substantiate that the development of a complex microenvironment using fibroblast profibrotic ECM (fECM) and the addition of TGFβ-1 allows us to recreate the fibrotic niche *in vitro* and can be considered as a feasible model for simulation fibrotic process *in vitro* for research and translational studies.

### Materials and methods Cell lines

The used cell cultures were obtained within the frame of Lomonosov MSU Project "Noah’s Ark". Cell lines of primary MSCs from human adipose tissue, human dermal fibroblasts, and lung fibroblasts were obtained from the Biobank of the Institute for Regenerative Medicine, Medical Research and Educational Center, Lomonosov MSU (ID: MSU_MSC, MSU_FB, https://human.depo.msu.ru). MSC cell lines were grown using AdvanceSTEM™ nutrient medium (Cytiva, USA) supplemented with 10% AdvanceSTEM™ Supplement (Cytiva, USA) and 1% penicillin and streptomycin (Gibco, USA). Skin fibroblasts and lung fibroblasts were cultivated in DMEM with 1g/L glucose (Gibco, USA) supplemented with 10% fetal bovine serum (FBS, Cytiva, USA) and 1% penicillin and streptomycin (Gibco, USA). All lines were cultured under standard conditions 5% CO2 and 37°C.

### Preparation of profibrogenic decellularized extracellular matrix (fECM)

The decellularized matrix was prepared as described previously [16]. Briefly, to form a cell layer, fibroblasts were planted at a concentration of 60 000 cells/ml on culture plastic (Corning) in DMEM with 1g/L glucose medium, with the addition of 1% antibiotic-antimycotic, 10% FBS and 50 µg/ml ascorbic acid (Sigma, USA) for two weeks. The medium was changed every two days. The resulting cell layer was decellularized using the detergent CHAPS (0.5% for 3 min) and DNase type I (Sci-Store, Russia, 50 U/ml, 30 min, 37°C), and then subsequently used as a profibrogenic extracellular matrix (fECM).

### Analysis of fECM structure and composition

Samples of fibroblast cell sheets and decellularized ECM (dECM) were fixed in 10% neutral formalin, then they were washed in distilled water and dehydrated in alcohols of ascending concentration, a mixture of alcohol-acetone and then pure acetone. Samples were dried at the critical point using Quorum K350 (Quorum gala instrument gmbh, Germany) or HCP-2 (Hitachi, Japan) was used. The samples were mounted on a special aluminum table with conductive carbon glue, sprayed with gold or platinum-palladium alloy in the spraying unit Quorum Q150TS (Quorum gala instrument gmbh, Germany) or IB-3 Ion Coater (EIKO, Japan) and observed with scanning electron microscope S 3400N (Hitachi, Japan). Type I collagen, fibronectin and EDA-fibronectin in the deposited ECM were evaluated by immunohistochemical labeling.

### Modeling of profibrotic conditions

To mimic profibrotic conditions *in vitro*, MSCs were seeded onto the resulting fibroblast fECM in serum-free DMEM supplemented with 5 ng/ml TGFβ-1 (Cell Signaling). After 4 days, the conditioned culture medium was collected for further analysis by ELISA. The cells were washed from the remnants of the culture medium with PBS (PanEco, Russia). For further PCR analysis, cells were lysed in QIAzol Lysis Reagent (79306, Qiagen) or RLT (79216, Qiagen). For immunocytochemical analysis, samples were fixed in 4% neutral formalin (AppliChem Panreac).

### Immunocytochemical analysis

Fixed samples were permeabilized using 0.2% Triton X-100 solution. Nonspecific binding of second antibodies was then blocked for 1 hour using normal 10% goat serum (Abcam) prepared in 1% BSA (Thermo Fisher Scientific). Antibodies to αSMA (ab5694), collagen I (ab34710), fibronectin (ab2413) and EDA-fibronectin (ab11575) (all - Abcam, UK) were used to label the studied targets in cells. Antibody detection was performed using second antibodies conjugated to a fluorescent label for 1 hour at room temperature in the dark: goat anti rabbit (Invitrogen, A11034 or A11037). To detect fibrillar actin, cells were also incubated with phalloidin conjugated to Alexa 594 (Molecular probe, A12381). Nuclei were labeled with DAPI (Sigma). Microscopic examination was performed on a Leica DMi8 microscope equipped with a Leica DFC 7000 T camera (Leica Microsystems GmbH), using representative fields of view to take photographs.

### Linked immunosorbent assay

The quantity of hepatocyte growth factor (HGF), basic fibroblast growth factor (FGF2), secreted protein acidic and rich in cysteine (SPARC, or osteonectin), interleukins 8 (IL-8) and 10 (IL-10), fibulin type 2 (fibulin-2) in conditioned medium samples was determined using appropriate commercial ELISA kits (R&D Systems, USA, for HGF, FGF2, SPARС, IL-8, IL-10 and Cloud-Clone Corp., USA for fibulin-2) according to the manufacturer’s instructions.

### Estimation of mRNA level of collagen I, EDA-FN and αSMA by real-time PCR

To isolate RNA (>200 bp) from cell lysates, a commercial kit RNeasy Mini Kit (74106, Qiagen) was used. RNA was isolated according to the protocol included with the kit. RNA concentration after purification was measured using a Nanodrop spectrophotometer (Thermo Scientific). Reverse transcription was performed using the MMLV RT kit (Evrogen). Real-time PCR was performed using qPCRmix-HS SYBR+LowROX (Evrogen) and primers presented in Table 1.

**Table 1.**
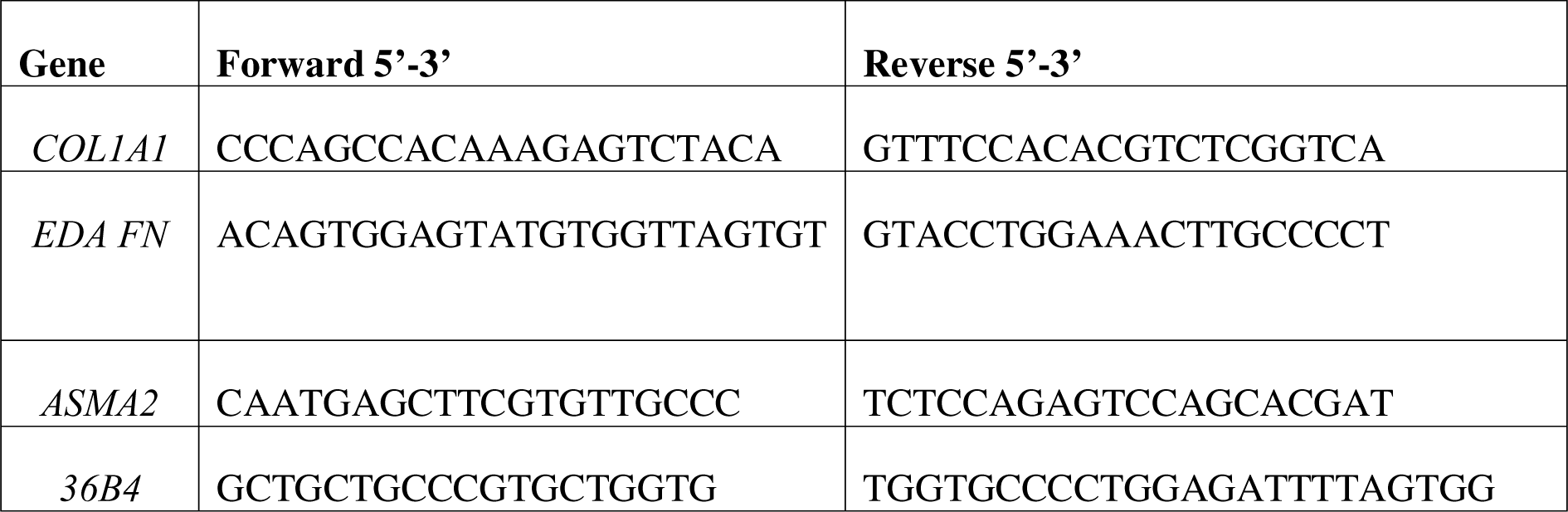
Primers sequences.

RT-PCR was performed on a QuantStudio 5 Real-Time PCR System (Applied Biosystems, Thermo Fisher Scientific) using a commercial protocol. The expression level of target mRNAs were calculated relative to the level of control mRNAs using the ΔCT calculation method.

### Immunoblotting

Culturing cells were lysed on ice in 2x Laemmli sample buffer (20 mM dithiothreitol, 6% SDS, 0.25 M Tris pH=6.8, 10% glycerol, 4% β-mercaptoethanol, 10 mM bromophenol blue). Protein level was measured using the DC Protein Assay kit (Bio-Rad, USA) according to the manufacturer’s instructions. Cell lysate proteins were separated by denaturing electrophoresis in the presence of 20% SDS (Sigma). PageRuler™ Plus (Invitrogen), a standard commercial prestained protein mixture of known molecular weight, was used to determine protein molecular weights. At the end of electrophoresis, proteins were transferred by wet blotting from the gel onto an Amersham Hybond-P PVDF membrane (GE Healthcare, UK) at a voltage of 100 V for 60 minutes. Nonspecific antibody binding was blocked in TBS/Tween containing 5% skim milk for 40 minutes at room temperature. Immunoblotting was performed using unlabeled monoclonal antibodies that specifically bind to GAPDH (Cell Signaling Technology, USA, 1:1000), αSMA (Abcam, 1:500), FAPa (Cell Signalling, 1:1000) overnight at +4°C and subsequent incubation with secondary antibodies 1:3000 (goat anti-rabbit IgG or rabbit anti-mouse IgG) conjugated with horseradish peroxidase (Sigma, USA) for 60 minutes at room temperature. Proteins bound to antibodies were visualized using a chemiluminescent substrate (ECL Pico or Dura, Pierce). A ChemiDoc UV screen (BioRad, USA) was used to expose the membranes. Chemiluminescence detection was carried out using the ChemiDoc Imaging System (BioRad, USA), and densitometric analysis of the resulting images was carried out using the ImageLab program.

### Estimation of fibrosis-associated microRNA (miRNA) content in MSCs using targeted PCR analysis

A commercial miRNeasy Mini Kit (217004, Qiagen) was used to isolate miRNA-enriched total RNA (>18 bp) from the obtained MSC lysates. RNA was isolated according to the protocol attached to the kit. RNA concentration after purification was measured on a Nanodrop spectrophotometer (Thermo Scientific). Reverse transcription was performed using miScript II RT Kit (Qiagen) with HiSpec buffer to obtain cDNA of mature miRNAs only. To assess changes in the profile of miRNAs associated with fibrosis, real-time PCR was performed using a commercial miScript miRNA PCR Arrays plate with miRNA-specific primers presorbed in the wells and a miScript SYBR Green PCR Kit (Qiagen) containing 2x QuantiTect SYBR Green PCR Master Mix and a universal miRNA primer. PCR was performed on a QuantStudio 5 Real-Time PCR System (Applied Biosystems, Thermo Fisher Scientific) in the mode according to the commercial protocol. Expression levels of target miRNAs were calculated relative to the levels of control miRNAs within the used array by the ΔCT method according to the manufacturer’s protocol. For this purpose, the content of tested miRNAs was normalized to the signal level obtained for SNORD61, SNORD95, SNORD96A, SNORD68, SNORD72, RNU6B/RNU6-2 cDNAs. After real-time detection PCR, the obtained data on the threshold cycle values for each gene were processed according to the formula: N = 2^(-ΔCt), ΔСt = Сt1-Сthk, where Ct1 is the value of the threshold cycle for the investigated gene, Cthk is the average values of threshold cycles for control miRNA genes, N is the result in relative units.

### Bioinformatic analysis

For bioinformatic analysis, miRNAs upregulated by more than 150% and downregulated by more than 50% were taken in MSCs cultured on fECM supplemented with 5 ng/ml TGFβ-1 (Cell Signaling) relative to MSCs cultured under standard conditions. The miRnet online service was used to search for targets. Experimentally validated and predicted miRNA targets were collected from miRTarBase v7.0, TarBase v7.0 and miRecords databases of the online service. After exporting the target genes from miRnet online service, they were annotated using David service.

### Statistical analysis

Experimental data are presented as 10 to 90 percentile, line at meridian. Statistical processing was conducted using GraphPad Prism 9.0 software (GraphPad Software, USA). Mann-Whitney nonparametric test was used to check the statistical significance in data between experimental and control groups. Differences were considered statistically significant when p<0.05.

## Results

### Profibrotic microenvironment leads to myofibroblast phenotype and expression profile in culturing MSCs

To mimic a profibrotic microenvironment, we added the well-known profibrotic factor TGFβ-1 to MSCs or cultured them for 4 days on ECM produced by human fibroblasts, which were cultured at high density in cell sheets with the addition of ascorbic acid to increase the efficiency of deposition of profibrotic ECM (fECM). Using scanning electron microscopy, we showed that after decellularization, fECM retains a three-dimensional structure with stacked fibers. Using immunohistochemical analysis, we showed that cell sheets from dermal fibroblasts cultured in DMEM with 10% FBS and ascorbic acid have their own network of matrix proteins, including collagen type I, fibronectin, EDA-FN, which is preserved after decellularization (Fig. 1).

**Figure 1.**
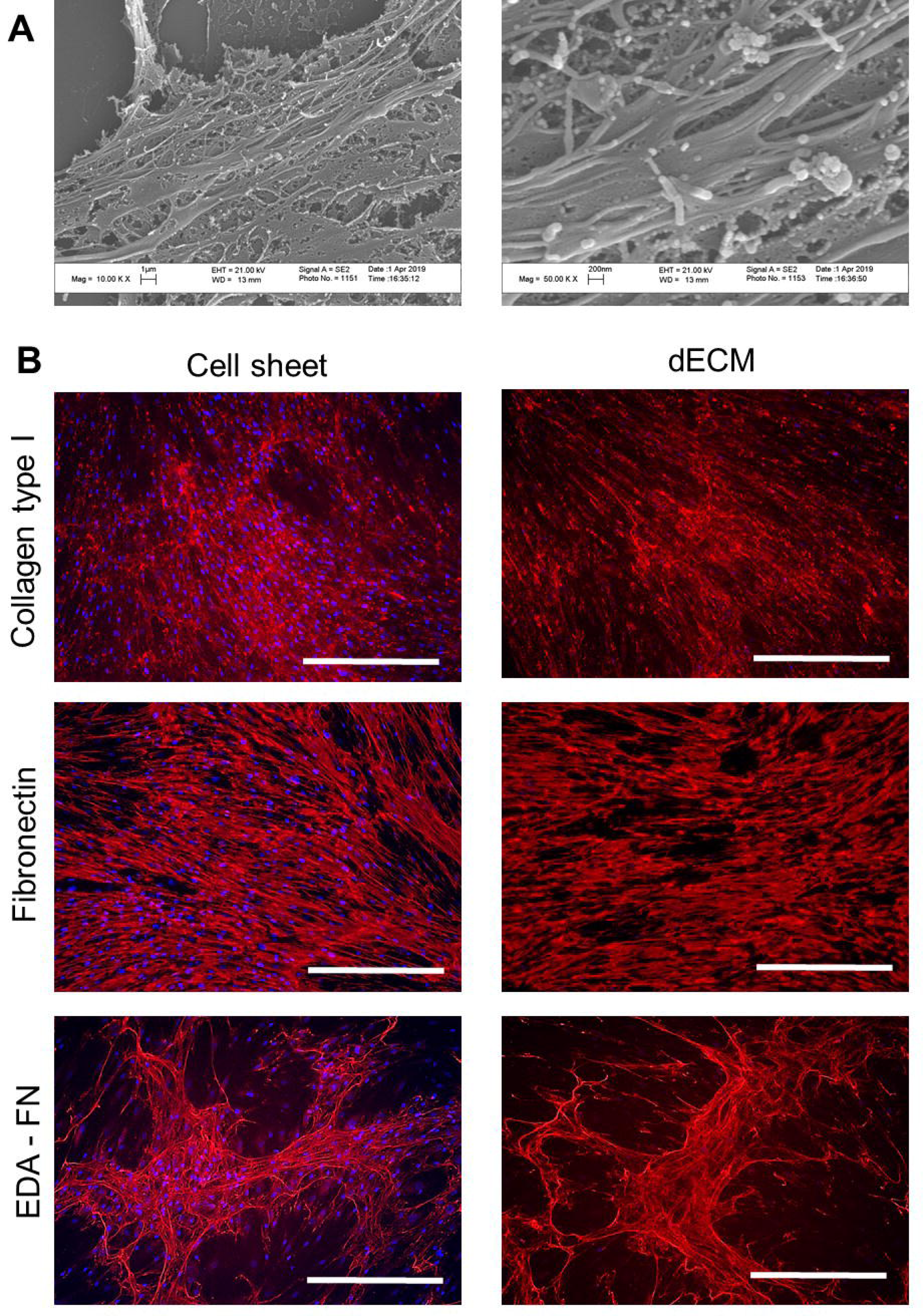
Characterization of profibrotic fECM. A. Scanning electron microscopy of dECM samples. Magnification: 1000x, 5000x. B. Analysis of ECM proteins in fibroblast cell sheets and dECM: type I collagen, fibronectin, EDA-FN (red) evaluated by immunohistochemistry. Fluorescent microscopy, nuclei stained with DAPI (blue), scale bar – 100 um.

Established culturing model was validated as profibrotic for various types of stromal cells, testing the effectiveness of stimulating the differentiation of cells into myofibroblasts when cultivated under such conditions. One of the main characteristics of the myofibroblast is an increased expression of the *ACTA2* gene, encoding αSMA, and the incorporation of αSMA into stress fibrils formed during myofibroblast differentiation. Thus, a mature myofibroblast acquires the ability to contract ECM realizing one of its crucial functions. Cultivation of MSCs under simulated profibrogenic conditions led to the appearance of αSMA+ stress fibrils after 4 days, which indicates the appearance of cells with a myofibroblast phenotype in MSC population (Fig. 2A). However, fECM itself did not cause the appearance of such cell phenotype.

**Figure 2.**
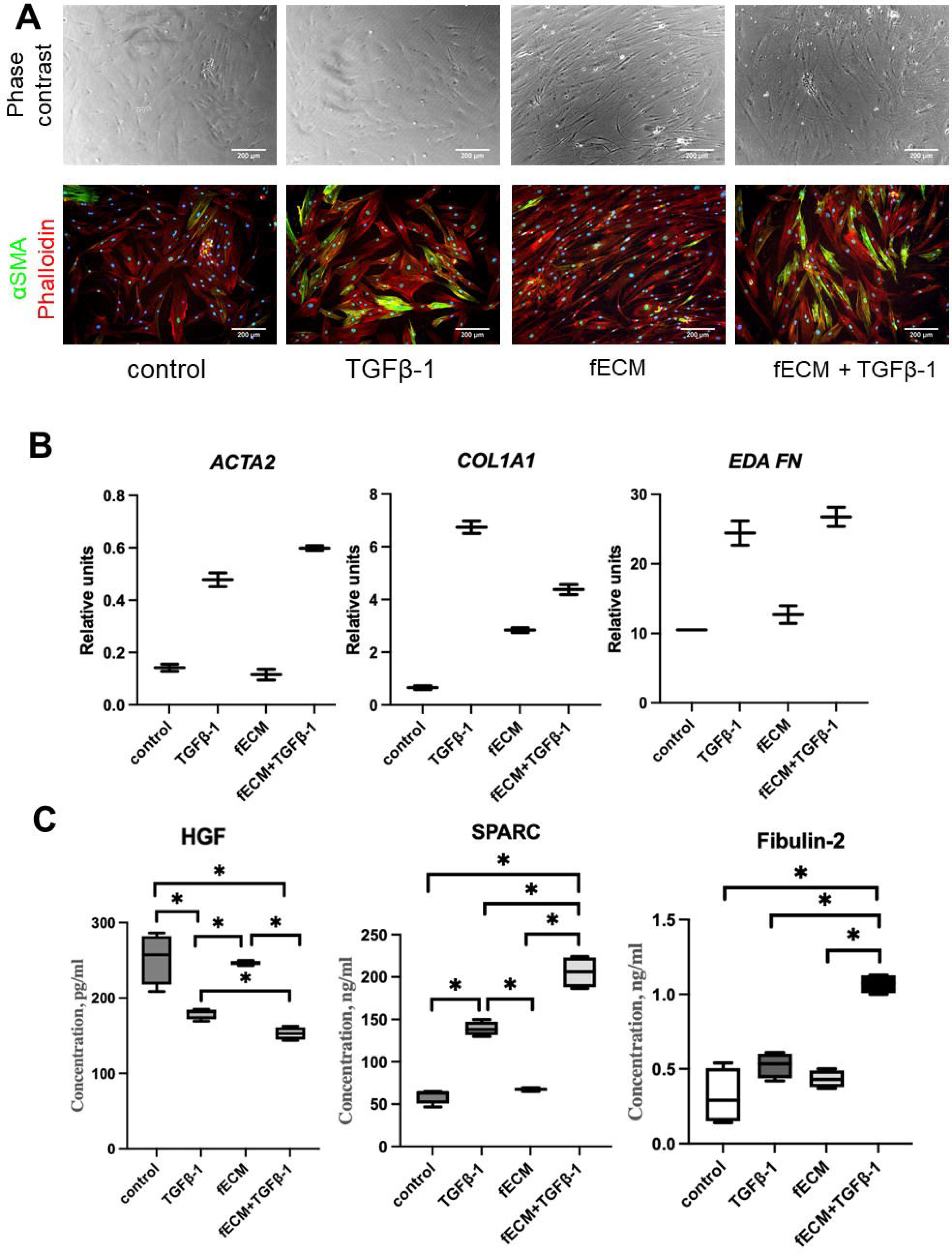
A. Morphology and expression of αSMA (green) in MSCs, cultivated in control condition, with 5 ng/ml TGFβ-1, on fECM or in complex profibrotic microenvironment (fECM+TGFβ-1) for 4 days. F-actin is labeled with phalloidin (red), nuclei are labeled with DAPI (blue). Phase contrast microscopy, fluorescent microscopy, scale bar - 200 um. B. Analysis of expression level of myofibroblast-associated genes *ACTA2*, *COL1A1*, *EDAFN* in MSCs, cultivated in control conditions, with 5 ng/ml TGFβ-1, on fECM or in complex profibrotic microenvironment (fECM+TGFβ-1) for 4 days. Data are presented as 10-90 percentile, line at median. n=2. C. The level of secreted soluble proteins HGF, SPARC, fibulin-2 in conditioned media collected from MSCs cultivated in different conditions for 4 days measured by ELISA. Data are presented as 10-90 percentile, line at median. n=4. p<0.05 (*).

We observed an increased expression level of the genes for profibrotic factors (*ACTA2*, *COL1A1*, *EDA-FN*) in MSCs induced by TGFβ-1 exposure for 4 days. Modeling the profibrotic microenvironment by culturing MSCs on fECM and in the presence of TGFβ-1 led to a greater increase in the expression of these genes compared to TGFβ-1 alone (Fig.2B). Wherein, fECM alone did not make a significant contribution to changes in expression compared to the control.

### Profibrotic conditions changes the secretion profile of MSCs

Considering the important role of soluble factors secreted by MSCs in the implementation of their antifibrotic effects, we analyzed changes in the content of several key pro- and antifibrotic protein mediators in the conditioned medium of MSCs cultured under various conditions. It was shown that under the influence of profibrotic stimuli, the secretory profile of MSCs changes in a coordinated manner: the cells secrete less HGF, which has an antifibrotic effect, and the level of secretion of one of the most important secretory regulators of TGFβ-signaling pathways and fibrosis of the protein SPARC and a representative of the secreted components of the fibrotic ECM protein fibulin-2 increases significantly (Fig. 2C). The maximum difference with the control was observed in the model of a profibrotic environment under the influence of a combination of fECM and TGFβ-1, while TGFβ-1 itself induced changes to a lesser extent, and fECM did not lead to significant differences. At the same time, the production of factors such as FGF2, IL-8, IL-10 by cells remained virtually unchanged (data not shown).

### Analysis of microRNA expression profile in MSCs cultured under profibrotic conditions

Analysis of the content of fibrosis-related miRNAs in MSCs, including their alteration under profibrotic conditions (fECM+TGFβ-1) using a targeted PCR assay showed that 74 miRNAs out of 84 miRNAs included in the analysis were expressed in the cells. Among them, 6 miRNAs (miR-132-3p, miR-133a-3p, miR-216a-5p, miR-215-5p, miR-203a-3p, miR-5011-5p) were expressed only in cells cultured under profibrotic conditions, and 2 miRNAs (miR-141-3p, miR-211-5p) only under standard conditions.

Under the influence of profibrotic stimuli, the expression of 49 miRNAs increased more than 1.5-fold in MSCs (Table S1). They had 9802 predicted targets identified by bioinformatic analysis, 9795 of which were unique. Clustering of these targets showed that a significant proportion of them are related to ECM production and remodeling, morphogenesis, regulation of intracellular signaling pathways involved in epithelial-mesenchymal transition, cell differentiation, cell cycle, and catabolic processes in cells. However, the analysis of common targets for these miRNAs (Fig. S1) does not allow us to clearly define the shift of the expression profile towards pro- or antifibrotic: among the target genes there are mediators that are multidirectional in their effects with regard to fibrosis regulation. Analyses for key fibrosis-associated terms confirm these findings.

Under profibrogenic stimuli, only 3 miRNAs were reduced in MSCs by almost 50% or more (Table S1). These miRNAs had 1362 unique predicted targets identified by bioinformatic analysis. Clustering of these targets showed that most of them are related to the regulation of protein synthesis, epithelial-mesenchymal transition and factors involved in angiogenesis. No common target genes regulated by all these miRNAs were identified. The groups of target genes identified by key terms associated with fibrosis are presented in the Supplement (Fig. S1). Interestingly, some of them overlap with miRNA targets whose expression is significantly upregulated under profibrotic conditions.

### Profibrotic microenvironment increases FAPa expression in human fibroblasts and their differentiation into myofibroblasts

The developed model of profibrotic microenvironment demonstrated the ability to promote the direct differentiation of MSCs into myofibroblasts. To further validate our model we used other stromal cell types, human dermal fibroblasts and human lung fibroblasts, and evaluated their behavior under profibrotic conditions. We found that culturing fibroblasts on fECM did not lead to the appearance of cells with a myofibroblast phenotype with manifest αSMA+ stress fibrils (Fig. 3A). On the contrary, supplementation of TGFβ-1 led to the differentiation of myofibroblasts after 4 days. The combined effect of fECM and TGFβ-1 also promoted the differentiation of the majority of fibroblasts into myofibroblasts.

**Figure 3.**
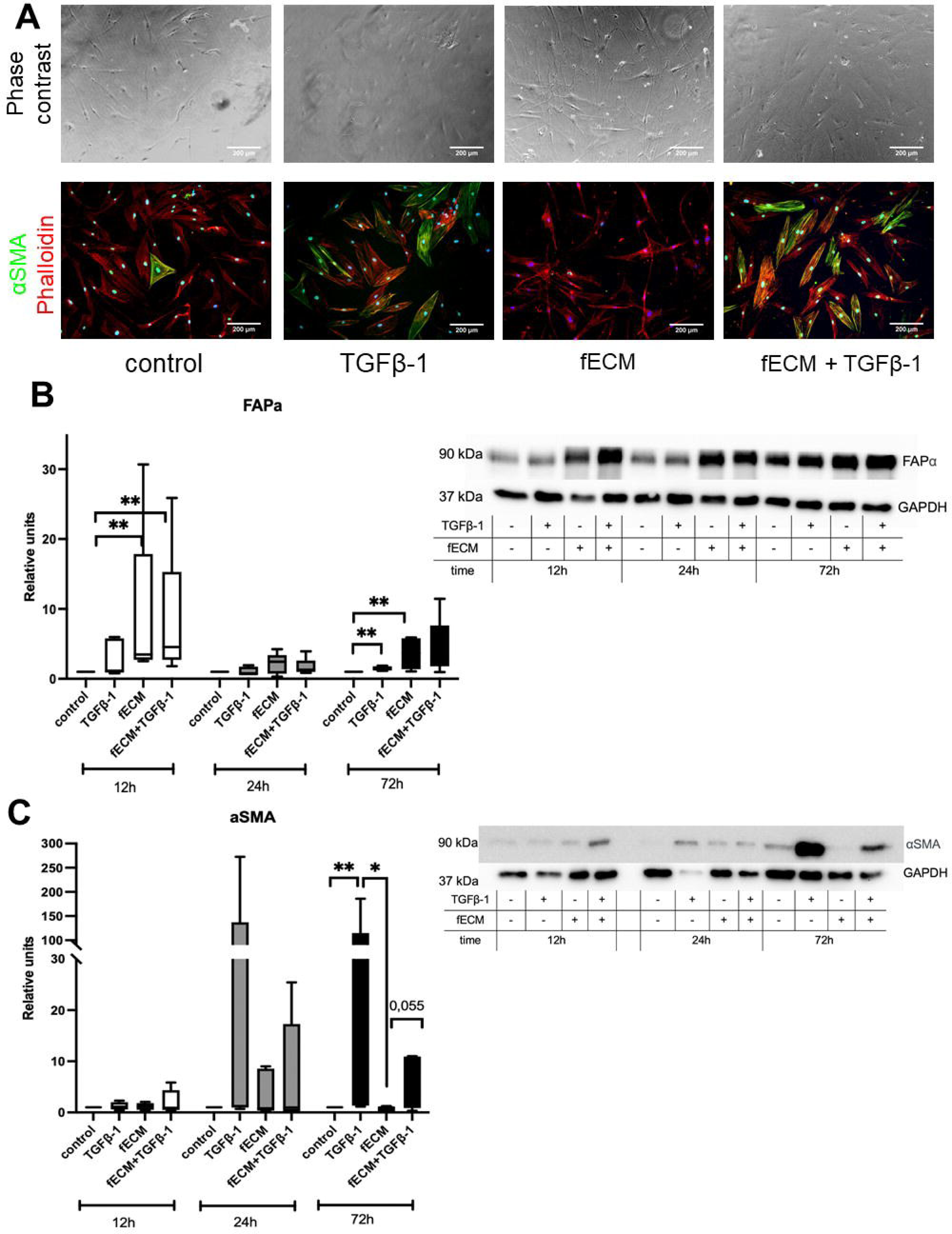
A. Morphology and expression of αSMA (green) in human dermal fibroblasts, cultivated in control condition, with 5 ng/ml TGFβ-1, on fECM or in complex profibrotic microenvironment (fECM+TGFβ-1) for 4 days. F-actin is labeled with phalloidin (red), nuclei are labeled with DAPI (blue). Phase contrast microscopy, fluorescent microscopy, scale bar - 200 um. B. Representative immunoblot image and calculated level of FAPa in human lung fibroblasts lysates, cultivated in control condition, with 5 ng/ml TGFβ-1, on fECM or in complex profibrotic microenvironment (fECM+TGFβ-1) for 12, 24 or 72 h. Data are presented as 10-90 percentile, line at median. n=5. p<0.005 (**). C. Representative immunoblot image and calculated level of αSMA in human lung fibroblasts lysates, cultivated in control condition, with 5 ng/ml TGFβ-1, on fECM or in complex profibrotic microenvironment (fECM+TGFβ- 1) for 12, 24 or 72 h. Data are presented as 10-90 percentile, line at median. n=5. p<0.05 (*), p<0.005 (**).

We analyzed the αSMA and FAPa level changes in human fibroblasts cell lines after seeding on fECM and exposure to TGFβ-1 after 12, 24 and 72 hours. Immunoblotting data showed that fECM led to increased FAPa level after 12 hours (Fig.3B). Moreover, fECM itself did not lead to elevation of αSMA level in fibroblasts after 12, 24 and 72 hours (Fig. 3C). With the combined exposure of fECM and TGFβ-1, the increase in αSMA expression was less than with the action of TGFβ-1 alone; however, the level of FAPa remained elevated on fECM compared to cells cultured on plastic and after 72 hours of cultivation.

## Discussion

Modeling fibrosis is an important issue both for fundamental science to understand the mechanisms of tissue repair and for translational studies to search for promising targets and methods to stop and reverse the process of fibrosis. Existing animal models have a number of significant limitations. For example, a well-established model of bleomycin-induced pulmonary fibrosis in mice does not lead to the formation of a pathologically self-sustaining process of lung tissue replacement by connective tissue, so researchers observe complete recovery of lung tissue after several months [17]. Moreover, *in vivo* system is complex, making it challenging to identify the role of individual molecules and cell types. Finally, the animal body is not able to fully reproduce the processes characteristic of human cells and tissues. In this regard, there is a urgent need for the relevant *in vitro* cellular models to investigate the fibrotic processes. On the one hand, they allow simplifying the system, making it possible for researchers to make unambiguous conclusions about cellular and molecular targets of interest, subject to the study of human cellular material. On the other hand, such models should, to the maximum extent possible, reproduce the complexity of the specific microenvironment *in vivo*. The characteristics include tissue architecture, the presence of intercellular contacts, including between different cell types, paracrine interactions, contact with a tissue-specific ECM [18].

ECM is an integral part of the tissue, participating in the regulation of homeostasis in the functional and damaged tissue [19]. In particular, ECM takes part in maintaining the state of stem cells and vice versa, the release of stem cells from the dormant state and the regulation of the differentiation of progenitor cells. Individual ECM proteins have long been used by different authors to elucidate their functions in the mentioned processes [20, 21]. However, the requirements for recreating the complexity of the system lead to the emergence of models in which the authors combine several types of ECM components [22] or three-dimensional models to recreate the three-dimensional architecture of tissues [23]. For maximum accuracy in recreating the profibrotic matrix, we have developed a method for its production by human fibroblasts with the addition of ascorbic acid, which is a cofactor of prolyl hydroxylase required for the maturation of collagen fibers. In addition, we have shown that the matrix obtained by this approach (fECM) contains a large amount of EDA-FN, known to be involved to the activation of fibroblast differentiation into myofibroblasts along with the rigidity of the ECM and the action of active TGFβ-1[24]. In addition to preserving the multicomponent chemical composition of the resulting matrix, it retains certain architecture providing the matrix scaffold for the differentiating stromal cells.

MSCs have the ability to differentiate under the influence of inducers in the direction of osteocytes, adipocytes, chondrocytes, myocytes [25], and in addition, they can be also one of the sources of myofibroblasts in the tissues [26]. We have shown that stromal cells cultured on fECM changed their morphology, becoming less spread out, more elongated, and retaining a 3D structure to a greater extent compared to cells cultured under standard conditions on plastic. However, culturing MSCs on fECM itself did not lead to the significant stimulation of myofibroblast marker genes expression (*ACTA2*, *COL1A1*, *EDAFN*), the appearance of αSMA+ stress fibrils, or changes in the level of secretion of HGF, SPARC, or fibulin-2. This result correlates with earlier observations, reporting that EDA-FN itself did not stimulate myofibroblast phenotype but is essential for TGFβ-1 stimulation [13]. We have previously shown that culturing stromal cells on ECM obtained from both fibroblasts isolated from the lungs of healthy mice (nECM) and the lungs of mice with bleomycin-induced fibrosis (blECM) did not lead to significant differentiation of fibroblasts into myofibroblasts [27]. However, a significant difference was observed with TGFβ-1, which is considered a major profibrotic paracrine factor and has been used by many researchers *in vitro* to induce myofibroblast differentiation. Induction with TGFβ-1 on the fECM did not lead to the appearance of myofibroblasts, while the number of αSMA+ positive cells and the level of αSMA increased 2-fold compared to the control with TGFβ-1 stimulation of fibroblasts on the blECM [27].

In the developed model of profibrotic environment, we also noted the potentiating effect of fECM on TGFβ-1-induced MSC differentiation into myofibroblasts. The effect of a combination of fECM and TGFβ-1 on MSCs led to a more pronounced increase in the expression of *ACTA2*, *COL1A1*, *EDAFN*. There was a decrease in the secretion of one of the well-described antifibrotic soluble factors, HGF, greater than that observed with TGFβ-1 stimulation alone. In accordance with these changes, the secretion of SPARC, a secreted protein acidic and rich in cysteine, increased significantly, for which stimulation of the YAP/TAZ signaling cascade with further translocation of non-phosphorylated YAP into the nucleus and activation of target genes YAP, TAZ, CTGF and CYR61, and the formation of myofibroblasts was shown [28]. There was also an increase in the secretion of fibulin-2, an ECM protein that is required for the activation of TGFβ-1 signaling and mediates the development of fibrosis [29].

Importantly, we also observed the concerted response to profibrotic stimuli in miRNA expression changes in MSCs. In general, the bioinformatic analysis of predicted gene targets of differently expressed miRNAs indicated the significant shift to the transcriptomic pattern involved in the activation of TGFβ-1 signaling, ECM production and remodeling, stimulation of neovascularization and inflammation as well as epithelial-to-mesenchymal transition. Recently, we demonstrated the important contribution of selected miRNAs (miR-21, -29c, -129) secreted by MSCs to the antifibrotic effects of these cells on the model of pulmonary fibrosis [30] and for the regulation of fibroblast-to-myofibroblast differentiation in vitro [4]. The expression of these miRNAs was also upregulated in MSCs in the developed profibrotic microenvironment possibly reflecting the adaptive response of these cells to profibrotic signals.

Thus, our suggested approach allowed recreating the microenvironment which effectively stimulated the differentiation of MSCs into myofibroblasts, accompanying by the appearance of stress fibrils in αSMA+ cells, an increase in the expression of myofibroblast marker genes and the corresponding changes in cell secretome and miRNA expression pattern.

Currently, various cell types are considered as the source of myofibroblasts in tissue, including circulating fibrocytes, bone marrow progenitors, epithelial and endothelial cells. However, tissue resident fibroblasts are considered to be the main source of myofibroblasts [31]. Thus, we validate the developed model using human fibroblasts cultured on plastic or fECM with or without the addition of TGFβ-1. After 12, 24 and 72 hours, we assessed the levels of activated fibroblast protein (FAPa) and αSMA in the cells. FAPa is a membrane protease; this protein is transiently expressed in some mesenchymal tissues of the embryo during embryogenesis, but is almost undetectable in adult tissues. Exceptions to this include the formation of some types of cancer, wound healing, and the progression of fibrosis [32]. Based on the available data, FAPa is involved in the remodeling of adult tissues, since its knockout does not lead to developmental disorders in mice [33]. FAPa is often considered as one of the markers of tumor-associated fibroblasts, but accumulating evidence suggests that its involvement in pathological remodeling processes may be broader. The proteolytic activity of FAPa in fibroblasts of the invasive front during the development of pulmonary fibrosis may contribute to their invasion and progression of fibrosis. At the same time, the fibrotic focus formed by a dense core of terminally differentiated αSMA myofibroblasts and the hard extracellular matrix they deposit due to the rigidity and composition of the ECM can induce the activation of new fibroblasts along its periphery by turning on the expression of FAPa [11, 34]. In our model, culturing fibroblasts on fECM led to an increase in FAPa levels both with and without the addition of TGFβ-1. At the same time, we observed an increase in the level of αSMA in cells induced by TGFβ-1 at 24 and 72 hours, and a significantly smaller increase in the level of αSMA at 72 hours in fibroblasts on the fECM under the influence of TGFβ-1. Cultivation on fECM did not lead to increased αSMA levels after 12, 24, and 72 hours. From the data obtained, we can conclude that fECM promotes the activation of fibroblasts and the acquisition of FAPa expression, which corresponds to the early initial stages of differentiation into myofibroblast. The combination of the action of fECM and TGFβ-1 allows us to obtain a later myofibroblast phenotype, which also expresses αSMA included in stress fibrils.

Taken together, the developed model of profibrotic microenvironment *in vitro* can be used to elucidate and clarify the mechanisms of fibrosis development in various tissues using tissue-specific cell lines as well as to test promising antifibrotic therapeutical agents and obtain a proof-of-concept of their action on cellular models.

## Funding

The study was supported by the Russian Science Foundation, grant No. 19-75-30007, https://rscf.ru/project/19-75-30007/ (dECM evaluation, validation of profibrotic model on MSCs) and grant No. 23-15-00198, https://rscf.ru/project/23-15-00198/ (validation of profibrotic model on fibroblasts).

## Acknowledgements

Equipment used for the study was purchased as a part of Lomonosov MSU Program of Development and Lomonosov MSU State Assignment.

## Notes

### Competing Interest Statement

The authors have declared no competing interest.

