## Supplementary figures and images for "Modeling the profibrotic microenvironment *in vitro*: model validation"

### Supplemental figure S1

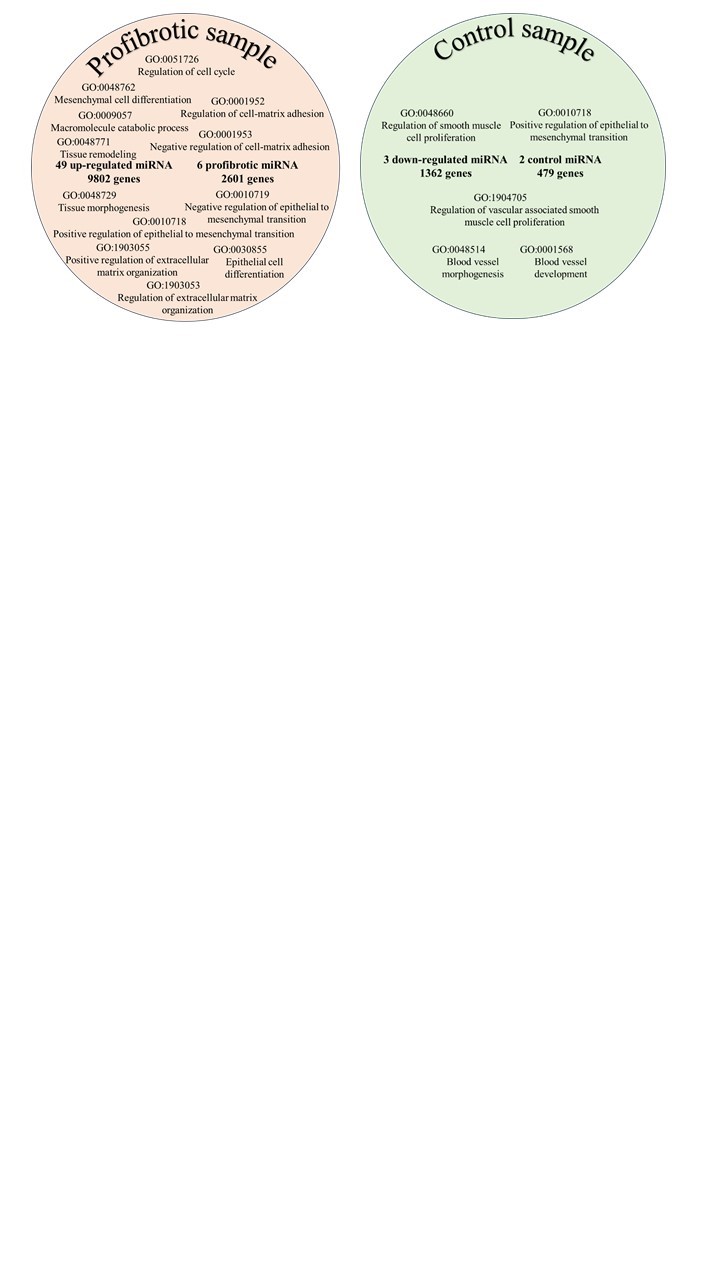

### Supplemental table s1

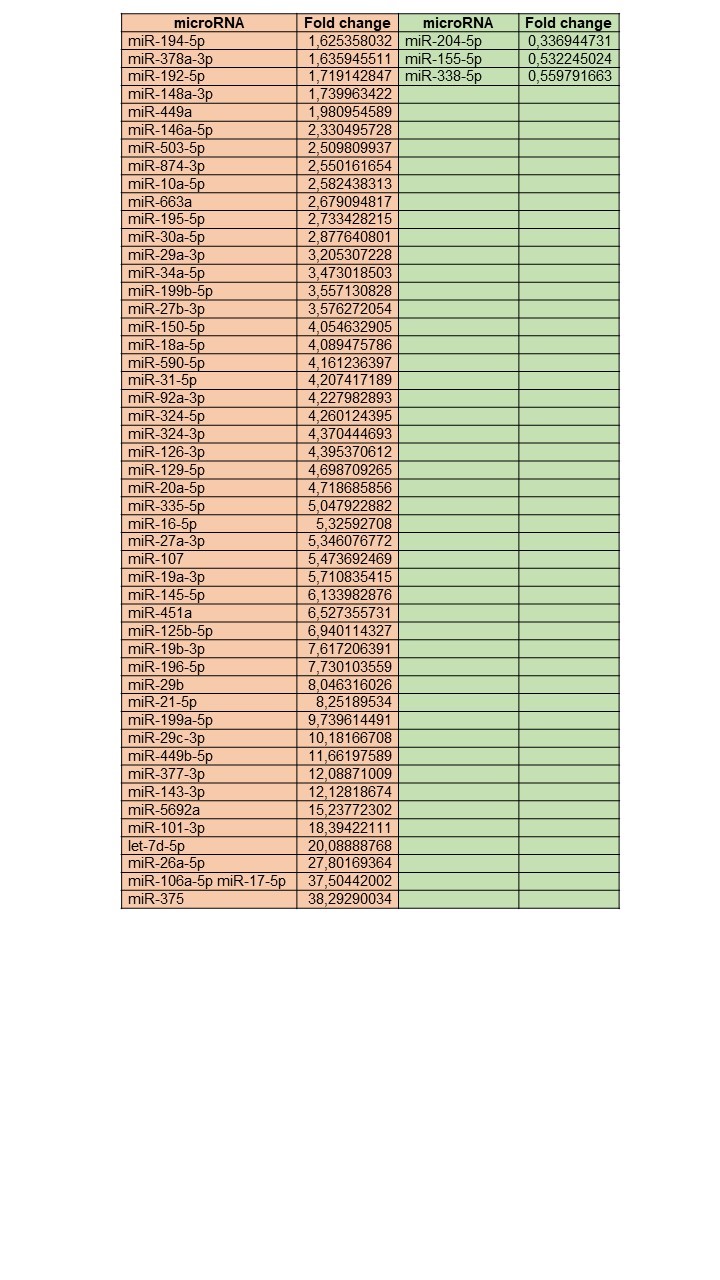
